# Beyond Peak Wavelength: Spectral Bandwidth of Blue and Red-Blue Laser Diodes (LDs) Reprograms Photosynthesis, Canopy Architecture, Senescence, and Whole-Plant Growth

**DOI:** 10.64898/2026.01.18.700227

**Authors:** Lie Li, Ryusei Sugita, Hiroyuki Togawa, Ichiro Terashima, Wataru Yamori

## Abstract

In indoor horticulture high planting densities often accelerate the senescence of lower leaves, increasing trimming frequency, reducing effective photosynthetic area, and raising labor costs. While spectral composition has been widely studied, the role of spectral bandwidth, particularly when peak wavelength is identical, remains poorly understood. Here, we used laser diodes (LDs) as narrow-band light sources to examine how spectral bandwidth influences photosynthesis, canopy architecture, leaf senescence, and plant growth under monochromatic blue light and combined red and blue (R+B) light in tobacco (*Nicotiana tabacum* L. ‘Wisconsin-38’), lettuce (*Lactuca sativa* L. ‘Red Fire’), and *Arabidopsis thaliana* (L.) Heynh. ‘Col-0’. Under monochromatic blue light, narrow-band LD blue (LD_B_) reduced CO□ assimilation rates and shoot dry weight compared with broad-band LED blue (LED_B_) across species. However, LED_B_ cultivation was accompanied by accelerated the senescence of lower leaves, a major limitation under dense planting. In contrast, plants grown under LD_B_ developed a more upright canopy architecture, characterized by larger leaf angles, which was associated with improved light penetration and delayed the senescence of lower leaves. When red and blue light were combined, narrow-band LD lighting (LD_R+B_) mitigated stress induced by 24-hour continuous illumination and promoted coordinated improvements in photosynthetic performance, leaf expansion, and canopy architecture. These integrated responses resulted in consistently higher shoot fresh weight and a healthier physiological state, as indicated by higher chlorophyll content and lower anthocyanin accumulation, compared with broad-band LED_R+B_. Together, our results demonstrate that spectral bandwidth, even when peak wavelength is held constant, is a critical parameter shaping plant growth strategies. Precise control of bandwidth enables partial decoupling of traits traditionally linked in the classic ‘sun’ or ‘shade’ leaf syndromes, allowing independent regulation of canopy architecture, leaf senescence, and plant growth. This study highlights spectral bandwidth as a powerful yet underutilized lever for optimizing canopy health and yield in indoor horticulture.

## 1. Introduction

Ensuring food security is an urgent global challenge driven by population growth, urbanization, and a declining agricultural workforce (UN, 2018; UN, 2022). Geopolitical instability and the increasing frequency of extreme climate events further threaten the stability of field-based production systems (Vogel et al., 2019; Behnassi and Haiba, 2022; Qu et al., 2023). In this context, indoor horticulture has emerged as a promising approach for sustainable food production, enabling year-round cultivation independent of climate, improving land-use efficiency through multi-layer systems, and allowing precise regulation of growth conditions (Levine et al., 2024; Furuta et al. 2025; Qiu et al. 2025; Takano et al. 2025). Because artificial lighting is both the primary driver of plant carbon gain and one of the major operational energy costs in plant factories, developing efficient lighting systems is essential to improve productivity and resource use efficiency (Kozai, 2013).

Light-emitting diodes (LEDs) have accelerated the development of “light recipes” by allowing control over wavelength composition, intensity, and photoperiod compared with conventional lamps. Numerous studies have shown that red (620–700 nm) and blue (400–500 nm) wavelengths are most effective for plant growth and can be combined to tune traits relevant to production, including photosynthesis, leaf expansion, and stomatal behaviors (Olle and Virsile, 2013; Ohtake et al. 2018; 2021; Saengtharatip et al., 2021; Van Delden et al., 2021). This effectiveness is due to the fact that these wavelengths are used not only serve as energy source for CO_2_ assimilation but also as signals to regulate plant development. Specifically, red and far-red light regulate development and acclimation largely through phytochromes (Strasser et al., 2010; Rao et al., 2011; Costigan et al., 2011), whereas blue light responses are mediated primarily by cryptochromes and phototropins, which control stomatal opening, chloroplast positioning, phototropism, and downstream metabolic and stress responses (Wang et al., 2008; Ando et al., 2022; Briggs and Christie, 2002; Landi et al., 2020).

However, a practical bottleneck in indoor systems is not only the instantaneous efficiency of photosynthesis but also canopy health under high planting densities. Dense canopies often intensify mutual shading and accelerate the senescence of lower leaves, increasing trimming frequency and labor while reducing the effective photosynthetic area that supports yield (Kim and Kubota, 2025; Tewolde et al., 2016). Whether the light environment can be strategically tuned to reshape canopy architecture and thereby mitigate these density-driven challenges remains unclear. Although spectral composition has been widely optimized, most LED-based studies have implicitly treated “blue” or “red” as single wavebands, despite the fact that typical LEDs emit across relatively broad spectral widths of around 50 nm (Li et al., 2025). Consequently, the specific impacts of spectral bandwidth remain insufficiently understood, even though it dictates how photons are distributed across pigment absorption features and photoreceptor action spectra. Variations in bandwidth inherently create distinct spectral distributions that may fundamentally alter the activation of light-harvesting of pigments and signaling pathways of photoreceptors, thereby reshaping leaf photosynthesis and canopy architecture, which may ultimately govern the trajectory of whole-plant growth and overall productivity.

Laser diodes (LDs) offer a distinctive opportunity to address this gap. In contrast to LEDs, LDs emit extremely narrow wavebands, often below 10 nm, enabling precise control over specific wavelengths in the delivered spectrum (Kasap, 2013; Li et al. 2025). In addition, laser light is coherent and highly directional (Fain and Milonni, 1987; Klimek-Kopyra et al., 2021), and LD-based systems can be compact, photon-efficient, and deliver light remotely with minimal heat load at the canopy (Hitz et al., 2012; Murase, 2015; Wierer et al., 2014). Despite these advantages, LDs remain underexplored as long-term horticultural light sources. Previous studies have mainly examined short-term or pre-illumination effects of laser irradiation on germination, early growth, and stress tolerance across diverse laser types and experimental settings (Klimek-Kopyra et al., 2020; Siyami et al., 2018; Nadimi et al., 2022; Ali et al., 2020; Cheng et al., 2025). Thus, how LDs influence whole-plant physiology, canopy architecture, and yield-related traits during sustained cultivation, particularly under demanding regimes such as continuous lighting, remains unclear.

Our recent study (Li et al., 2025) highlighted the potential application of LDs in indoor horticulture. We found that monochromatic red LDs (peak at 660 nm) substantially outperformed red LEDs (peak at 664 nm) in plant photosynthesis and growth. This study aims to determine whether the narrow spectral bandwidth of LDs can serve as a strategic tool to mitigate the physiological constraints of dense planting in indoor horticulture. First, we examined how monochromatic blue light sources differing in bandwidth (LD vs. LED) affect instantaneous photosynthetic responses, plant morphology, and acclimatory growth. Furthermore, we compared the effects of combined red and blue light from LDs and LEDs on plant morphology and growth, with a focus on how bandwidth influences canopy architecture and light interception. These experiments were conducted across three species, including tobacco (*Nicotiana tabacum*), lettuce (*Lactuca sativa*), and *Arabidopsis thaliana*.

## 2. Materials and Methods

In this study, three experiments were conducted using three plant species: tobacco (*Nicotiana tabacum* L. ‘Wisconsin-38’), *Arabidopsis thaliana* (L.) Heynh. ‘Col. 0’, and lettuce (*Lactuca sativa* L. ‘red fire’).

### 2.1 Experiment 1. Effects of different blue light spectra on gas exchange

#### 2.1.1 Plant materials, cultivation conditions, and gas exchange measurements under blue light treatments

Tobacco seeds were sown in a 1:1 vermiculite and peat mixture (Metro-Mix 350J, Hyponex, Japan) in black trays and thinned to one plant per pot post-germination. Seedlings were then grown in a growth chamber (LPH-411SPC, Nippon Medical & Chemical Instruments, Japan), under white fluorescent light providing a photosynthetic photon flux density (PPFD) of 150 μmol m□² s□¹. The environmental conditions were maintained as follows: a photoperiod of 10 h light/14 h dark, 25/22°C day/night temperatures, 60 ± 5% relative humidity, and CO□ concentration of 400 μmol mol^-1^. Fully expanded young leaves from four independent plants were selected for gas exchange measurements 25 days after sowing.

#### 2.1.2 Gas exchange measurements under blue light treatments

Gas exchange parameters, including net photosynthetic rate (P_n_) and stomatal conductance (g_s_), were measured using a portable gas exchange system (LI-6400XT with 6400-8 transparent leaf chamber, LI-COR Biosciences, Lincoln, NE, USA) (Yamori et al., 2016). Leaf chamber conditions were maintained at 25°C, 400 μmol mol□¹ CO□, and a leaf-to-air vapor pressure deficit (VPD) of 0.7–1.0 kPa. For this experiment, three blue light sources, LD 407 (peak wavelength 407 nm, waveband of 404∼411 nm, L405G1, Thorlabs, Inc., Japan), LED 450 (peak wavelength 450 nm, waveband of 431∼475 nm, 3LH-100DPS, Nippon Medical & Chemical Instruments, Japan), and LD 450 (peak wavelength 450 nm, waveband of 447∼454 nm, L450G1, Thorlabs, Inc., Japan) were used. Each light source provided a photon flux density (PPFD) of 150 μmol m□² s□¹. Spectral properties were measured using a compact spectrometer (CCS200/M, Thorlabs, Inc., Japan) and are presented in Figure 1A. Treatments were applied to four replicate leaves in a randomized sequence. Intrinsic water use efficiency (WUEi; mmol CO_2_ mol^-1^ H_2_O) was determined as the ratio of P_n_ to g_s_.

**Figure 1.**
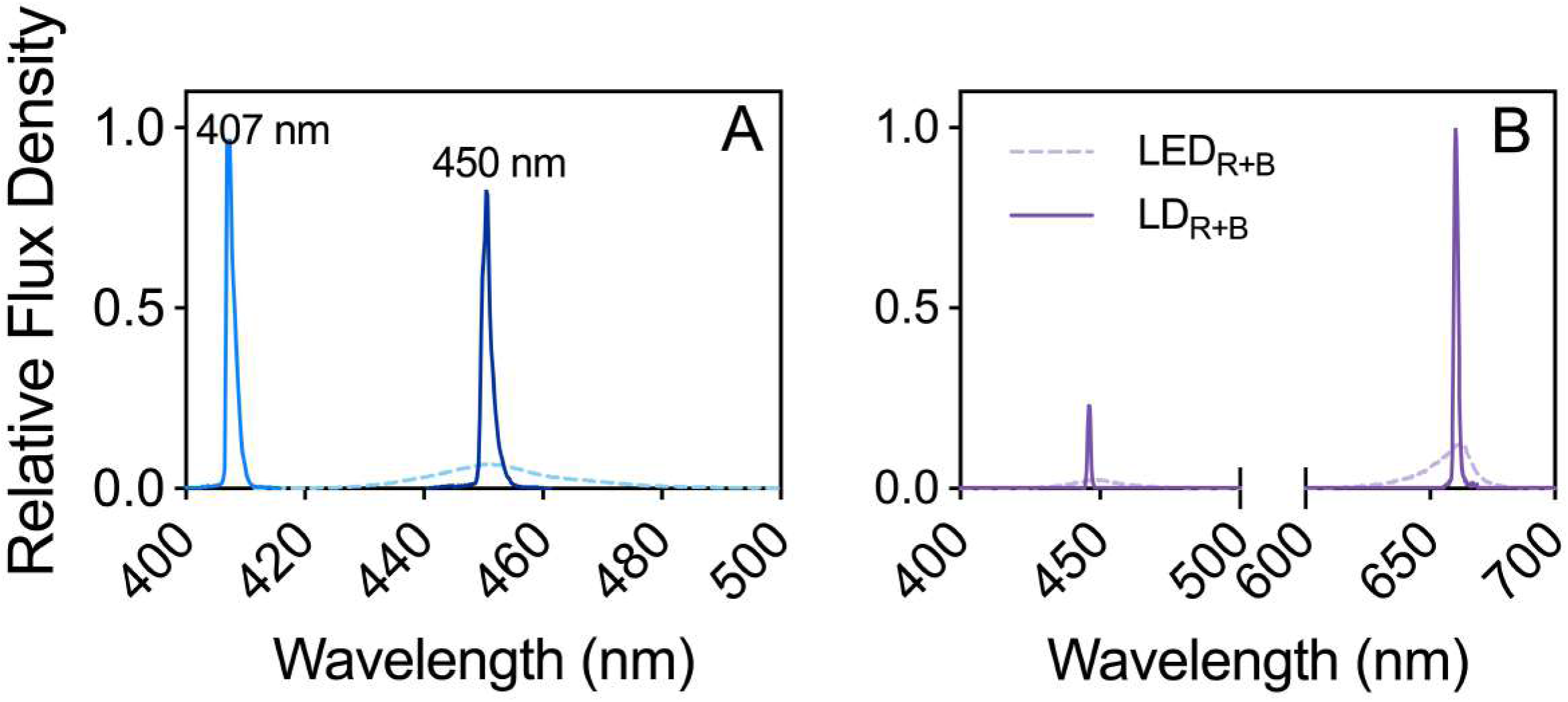
Spectral distributions of the light sources. The photosynthetic photon flux density (PPFD) for all treatments was maintained at 150 μmol·m□²·s□¹. (A) Spectra of the monochromatic blue LED (dashed line) and LD (solid) light sources. (B) Spectra of the combined red and blue (R+B) LED and LD light sources.

### 2.2. Experiment 2. Plant growth under monochromatic LED_B_ and LD_B_

To compare the effects of spectral bandwidth on plant growth, we subsequently selected blue light sources with identical peak wavelengths but different bandwidths, LED 450 (hereafter labeled as LED_B_) and LD 450 (hereafter LD_B_), for the following experiments.

#### 2.2.1. Plant materials, cultivation conditions, and light treatments

Tobacco seeds were sown and grown following the procedures as described in section 2.1.1. Arabidopsis seeds were sown in the same substrate as tobacco within plastic boxes and thinned to three seedlings per box. Lettuce seeds were sown in rockwool cubes saturated with distilled water and irrigated weekly with a 1/1000 strength nutrient solution (Hyponex 6-10-5, Hyponex Japan). Initial growth conditions for all seedlings were identical to those detailed in section 2.1.1. Light treatments specific to this experiment commenced when plants reached the three-true-leaf stage. At this point, seedlings were transferred to continuous 24-hour illumination for 12 days under either the LED_B_ or the LD_B_, both providing a PPFD of 150 μmol m□² s□¹. The environment conditions were controlled at 25 ± 1 °C, 60 ± 5% relative humidity, and 400 μmol mol^-1^ CO_2_. Four biological replicates were maintained for each species and light treatment combination. After the growth for 12 days, photographs were taken from both side and top of the plants. Subsequently, various growth indices were subsequently quantified for all biological replicates. Representative photographs for each species and treatment were presented.

#### 2.2.2. Plant growth and morphology analysis

After 12 days of exposure to either the LED_B_ or LD_B_ light treatments, growth and morphological parameters were assessed using four replicate plants per treatment for each plant species (n = 4). Leaf angle was defined as the angle between the leaf midrib and the horizontal plane; this was quantified from lateral-view images of fully expanded leaves using ImageJ software (version 13.0.6, National Institutes of Health, USA). Following leaf angle measurements, plants were harvested, and shoots were separated from roots using sterilized sharp scalpels. The total leaf area (cm^2^) of each plant was determined by scanning detached leaves (CanoScan LiDE 220, Canon Inc., Japan) and analyzing the images with ImageJ. Subsequently, shoots were oven-dried at 80°C for 72 hours to a constant weight to determine shoot dry weight (DW, mg). Leaf mass per area (LMA, mg cm□²) was then calculated as the ratio of shoot DW to the corresponding total leaf area.

#### 2.2.3. Pigment content analysis

After 12 days of growth under 24-hour continuous irradiation of either the LED_B_ or LD_B_ treatment, relative chlorophyll content (SPAD value; leaf area based) was estimated using a portable chlorophyll meter (SPAD-502PLUS, Konica Minolta, Japan). For each species (tobacco, lettuce, and Arabidopsis), measurements were taken from representative leaves at three canopy positions: upper, middle, and lower. On each selected leaf, three SPAD readings were taken from different points along the lamina, avoiding the midrib and major veins, and the average was recorded. Measurements were performed on four replicate plants per treatment for each plant species.

### 2.3 Experiment 3. Photosynthesis and plant growth under combined red and blue LED and LD

#### 2.3.1 Plant materials, cultivation conditions, and light treatments

Tobacco, lettuce, and Arabidopsis were used in this experiment. Seeds were sown, and seedlings were initially cultivated using the identical methods and controlled environmental conditions as described in section 2.2.1. Commencing at the three-true-leaf stage, seedlings were transferred to continuous 24-hour illumination for 15 days under either the combined red and blue LED (LED_R+B_) or the combined red and blue LD (LD_R+B_) source. Both light sources provided a PPFD of 150 μmol m^−2^ s^−1^, with a red to blue ratio of 4. The spectral distributions for these LED_R+B_ and LD_R+B_ sources were measured and presented in Figure 1B. Four biological replicates were maintained for each species and light treatment combination. After the 15-day growth period, photographs were taken from both side and top views, representative images were selected, and various growth indices were quantified.

#### 2.3.2 Chlorophyll fluorescence analysis

Photosystem II (PSII) efficiency was investigated in the youngest fully expanded leaves of all three species after 15 days of growth under either the LED_R+B_ or LD_R+B_ treatments. Measurements were performed using a pulse amplitude modulation fluorometer (Junior-PAM, Heinz Walz GmbH, Germany) by applying saturation pulses. After 30 min of dark treatment, either LED_R+B_ or LD_R+B_ light source was applied as actinic light at a PPFD of 150 μmol m^-2^ s^-1^. Once a steady-state fluorescence (F_s_’) was achieved, a saturating pulse was applied to determine the maximum fluorescence under actinic light (F_m_’). Four replicate leaves from separate plants of each species were measured under each actinic light condition. The quantum efficiency of PSII (Y(II)) was calculated using methods based on (Genty et al., 1989):

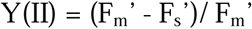

#### 2.3.3 Determination of spectrally weighted leaf absorptance and electron transport rate (ETR)

Leaf absorptance was measured on the same leaves used for chlorophyll fluorescence analysis. Leaf transmittance (T) and reflectance (R) in the range of 400-700 nm were determined using an integrating sphere equipped with a spectrometer (FLAME-Sl; Ocean Optics, Orlando, FL, USA). Light was provided by a halogen lamp (KL 1500 AL; SCHOTT, Mainz, Germany), via a fiber optic cable directed into the sphere’s top port, collimated using a collimator lens (Model MLS-60P, Moritex Corp., Japan). Before the sample measurement, the system was calibrated. A baseline was recorded by placing a 99% white reflectance standard at the bottom port while the light source was on, and this calibration was further validated against the known reference data provided by the manufacturer. A dark (0%) baseline was subsequently recorded with the light source turned off. To measure transmittance, a leaf disc was placed over the top port, with the white reflectance standard at the bottom port. To measure reflectance, the leaf disc was moved to the bottom port, and a black, non-reflective light trap was placed behind it. Each measurement was completed within 2 minutes. Leaf absorptance (A) was then calculated using the formula: A = 1 - R - T.

The spectrally weighted leaf absorptance (A_s_) was determined by integrating the measured leaf absorptance spectrum above weighted by the spectral distribution of LED_R+B_ and LD_R+B_ under which the plants were grown. Finally, ETR was calculated using the equation (Genty et al., 1989):

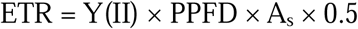

Where Y(II) is the measured quantum efficiency of PSII; PPFD is the photon flux density of actinic light used during chlorophyll florescence measurements; A_s_ is the calculated overall absorptance for the respective light sources; 0.5 represents the assumed fraction of absorbed PPFD distributed to PSII.

#### 2.3.4 Plant growth and morphology analysis

After 15 days of growth under either the LED_R+B_ or LD_R+B_ treatment, plants were harvested for analysis. Immediately upon harvest, shoot fresh weight (FW) was recorded. Shoot dry weight (DW), total leaf area, leaf mass per area, and leaf angle were determined as described in Section 2.2.2. The determination of leaf water content (%) was calculated as: [(FW−DW)/FW]×100. All measurements were performed on four replicate plants per species and treatment combination.

#### 2.3.4 Pigment content analysis

After 15 days of continuous 24-hour irradiation under either LED_R+B_ or LD_R+B_ treatments, leaf pigment content was assessed non-destructively. Relative chlorophyll content (SPAD values; leaf area-based) and anthocyanin content (ACI values; leaf area-based) were measured on the youngest fully expanded leaves using a portable chlorophyll meter (SPAD-502PLUS, Konica Minolta, Japan) and a portable anthocyanin meter (ACM-200 plus, Opti-Sciences, Inc., Hudson, NH, USA), respectively. For each species (tobacco, lettuce, and Arabidopsis), measurements were conducted on four replicate plants per treatment. On each selected leaf, three readings were taken from different positions across the leaf lamina, avoiding the midrib and major veins, and the average was recorded as a representative value.

### 2.4 Statistical analysis

Statistical analyses were performed following procedures similar to our previous work (Li et al., 2025). Specifically, to account for inter-individual plant variability in gas exchange data, a generalized linear mixed model (GLMM) was employed, followed by multiple mean comparisons using the Tukey□Kramer honest significant difference (HSD) test (*P* < 0.05) in R software 4.2.2 (R Development Core Team, 2021). Gas exchange data were visualized using box plots generated with GraphPad Prism 10 (GraphPad Software Inc., San Diego, CA, USA), with plot elements interpreted as described previously. For other parameters, data means were compared using independent t test (*ns* indicates not significant, \**P* < 0.05, \*\**P* < 0.01, and \*\*\**P* < 0.001) performed in SPSS 26.0 statistical software (SPSS Inc., Chicago, IL, USA). All graphs were plotted using GraphPad Prism 10.

## 3. Results

### 3.1 Plant responses to monochromatic blue LED and LD

#### 3.1.1 Effects of different blue light spectra on photosynthesis

Net photosynthetic rate differed significantly among blue light treatments (Fig. 2A). Tobacco leaves exposed to LD 407 and LED 450 exhibited 29.7% and 33.7% higher net photosynthetic rates, respectively, than those exposed to LD 450. No significant difference in net photosynthetic rate was detected between LD 407 and LED 450. In contrast, stomatal conductance did not differ significantly among treatments (Fig. 2B). Intrinsic water-use efficiency (WUEi) closely followed the pattern observed for net photosynthetic rate (Fig. 2C). Both LD 407 and LED 450 resulted in significantly higher WUEi values (28.2% and 34.3%, respectively) than LD 450, whereas no significant difference was observed between LD 407 and LED 450. These results indicate that differences in blue-light peak wavelength and bandwidth affected CO= assimilation efficiency, while having little effect on stomatal conductance.

**Figure 2.**
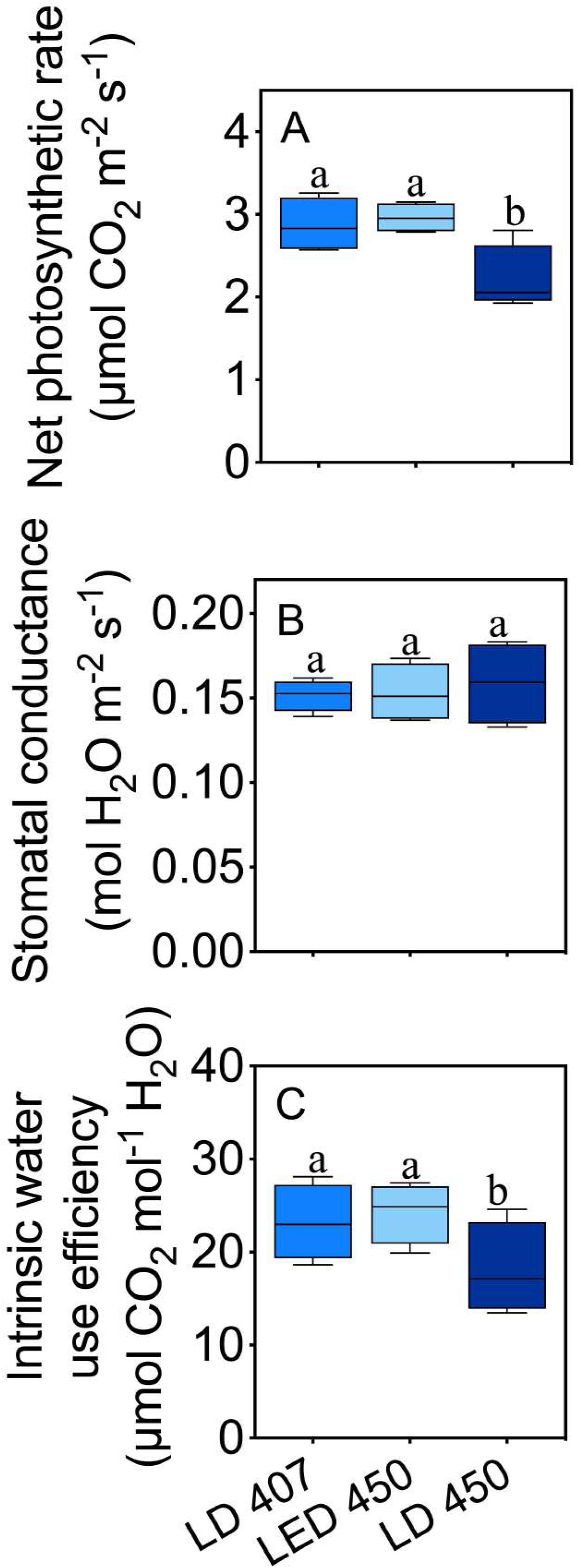
Gas exchange parameters of tobacco leaves under different monochromatic blue LED and LD light sources. Panels show (A) net photosynthetic rate, (B) stomatal conductance, and (C) intrinsic water use efficiency. Different letters above the columns indicate significant differences (*P* < 0.05, Tukey-Kramer HSD test). Data are presented as the mean ± SE (n = 4).

#### 3.1.2 Plant growth and morphology

Representative images of tobacco, lettuce, and Arabidopsis grown for 12 days under continuous monochromatic blue LED (LED_B_) or blue LD (LD_B_) irradiation are shown in Fig. 3A. Across all three species, plants grown under LD_B_ consistently exhibited a more upright canopy architecture, characterized by steeper leaf angles, whereas plants grown under LED_B_ showed a more horizontal leaf orientation. In addition, lettuce plants under LED_B_ accumulated visibly higher levels of anthocyanin, and Arabidopsis plants under LED_B_ initiated bolting and flowering earlier than those grown under LD_B_ (Fig. 3A). Quantitative analyses revealed a clear contrast between biomass accumulation and canopy architecture (Fig. 3B). Shoot dry weight was significantly greater under LED_B_ than under LD_B_, increasing by 47.8% in tobacco, 37.7% in lettuce, and 25.2% in Arabidopsis. In contrast, total leaf area did not differ significantly between light treatments in any species. Leaf mass per area (LMA) was consistently higher under LED_B_, with increases of 28.2% in tobacco, 27.6% in Arabidopsis, and 28.4% in lettuce relative to LD_B_. Conversely, leaf angle was markedly larger in plants grown under LD_B_. Mean leaf angles exceeded those under LED_B_ by 23.5° in tobacco, 13.7° in lettuce, and 17.4° in Arabidopsis. These results demonstrate that monochromatic blue LD induced pronounced changes in canopy structure that contrasted with the biomass accumulation observed under blue LED.

**Figure 3.**
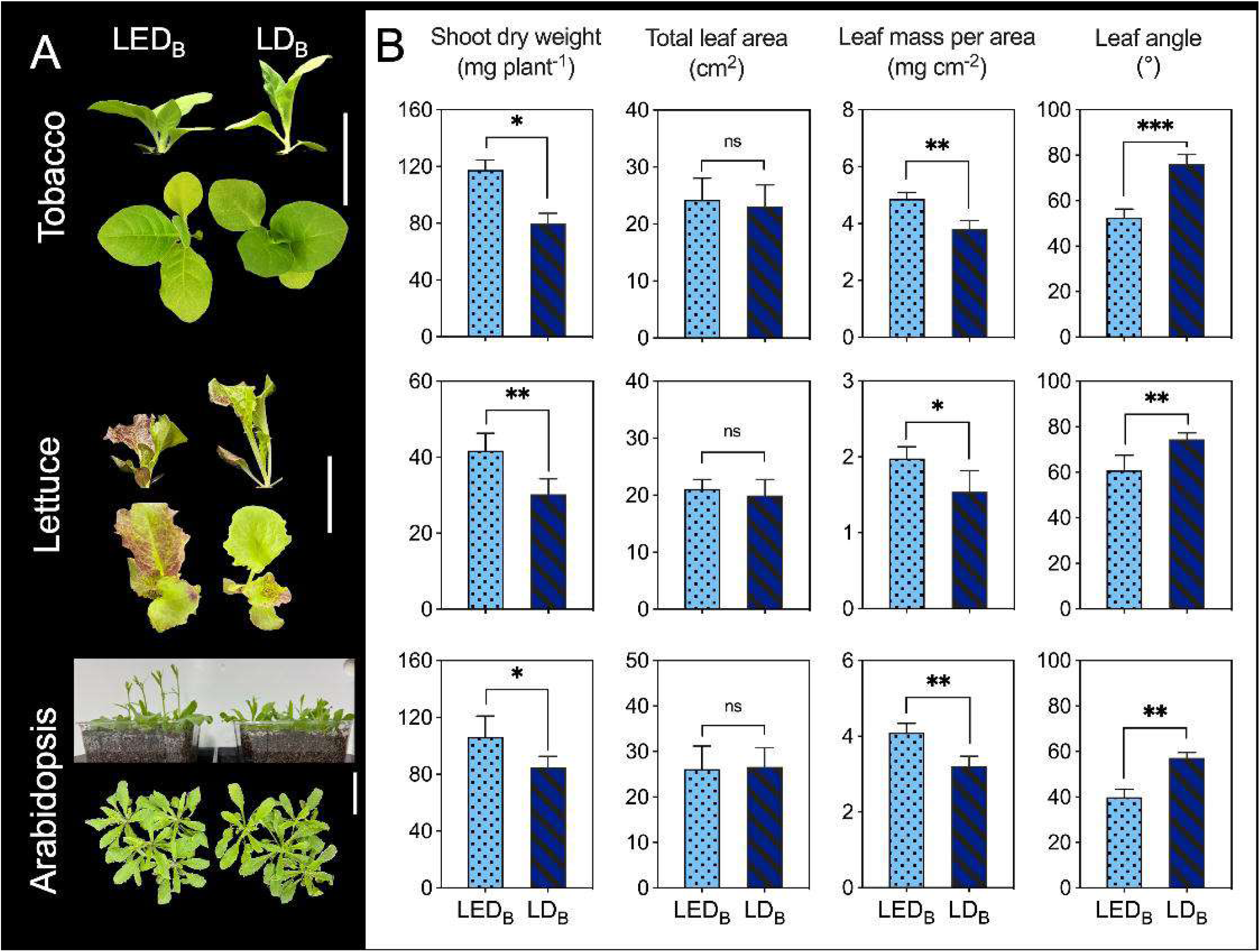
Growth and morphology of tobacco, lettuce, and Arabidopsis under continuous monochromatic blue light. Plants were grown for 12 days under either a monochromatic blue LED (LED_B_) or a blue LD (LD_B_) with a same peak at 450 nm at of PPFD of 150 μmol m□² s□¹. (A) Representative images of plants (side and top views). The white scale bar in each picture represents 5 cm. (B) Shoot dry weight, total leaf area, leaf mass per area, and leaf angle. Data are presented as the mean ± SE (n=4). Asterisks indicate significant differences between treatments (* *P* < 0.05, ** *P* < 0.01, *** *P* < 0.001; t test). ns indicates no significant difference.

#### 3.1.3. Pigment content

Relative chlorophyll content (SPAD value) was measured at three canopy positions (upper, middle, and lower leaves) for all species (Fig. 4). In tobacco, LD_B_-grown plants exhibited significantly higher SPAD values in lower leaves than LED_B_-grown plants, with an increase of 30.0%, whereas no significant differences were detected in upper or middle leaves (Fig. 4A). A similar pattern was observed in lettuce, where LD_B_ resulted in a 34.1% higher SPAD value in lower leaves compared with LED_B_, while SPAD values in upper and middle leaves did not differ significantly between treatments. In Arabidopsis, LD_B_-grown plants showed significantly higher SPAD values in both upper (+5.9%) and lower (+16.2%) leaves relative to LED_B_-grown plants. In contrast, SPAD values of middle leaves did not differ significantly between treatments in any species. Overall, these results indicate that monochromatic blue LD preferentially maintained chlorophyll content in lower canopy leaves compared with blue LED, despite lower whole-plant dry matter accumulation.

**Figure 4.**
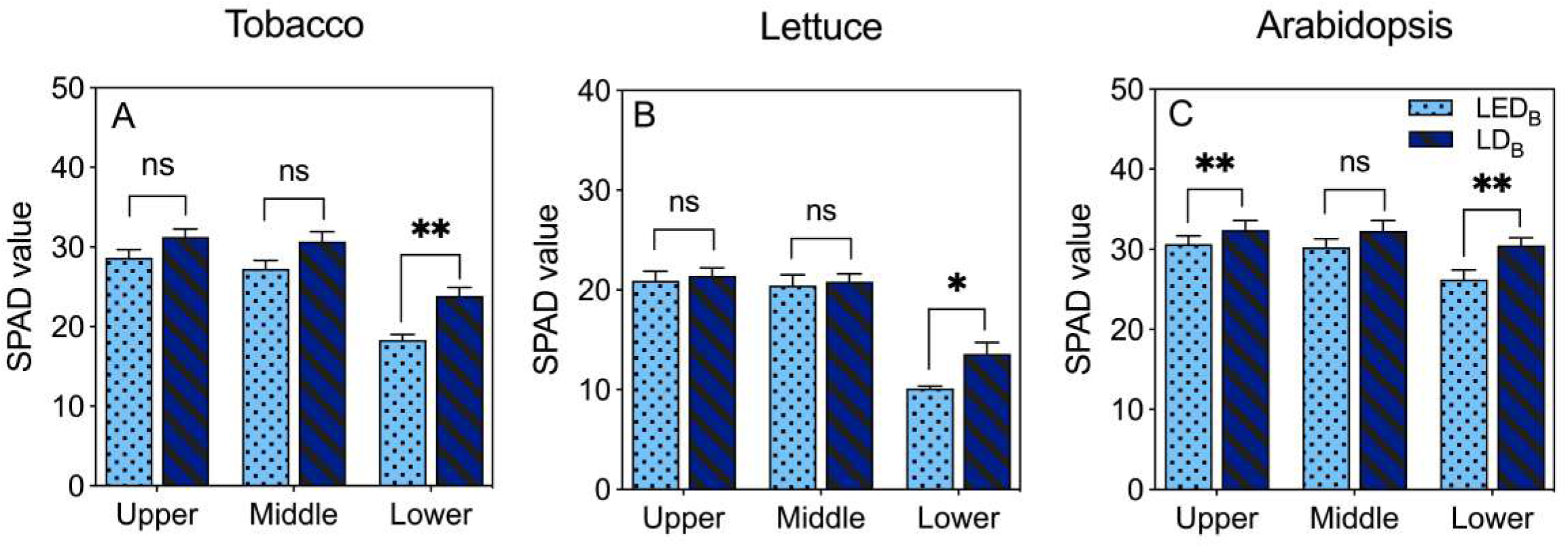
Relative chlorophyll content (SPAD value; leaf area based) in (A) Tobacco, (B) Lettuce, and (C) Arabidopsis leaves at different canopy positions. Plants were grown for 12 days under continuous light from either a monochromatic blue LED (LED_B_) or a blue LD (LD_B_), both with an identical peak wavelength at 450 nm and a PPFD of 150 μmol·m□²·s□¹. Measurements were taken from upper, middle, and lower leaves. Data are presented as the mean ± SE (n = 4). Asterisks indicate significant differences between treatments (* *P* < 0.05, ** *P* < 0.01; *t* test). ns indicates no significant difference.

### **3.2.** Plant responses to combined red and blue LED and LD

#### 3.2.1 Leaf absorptance spectra, spectrally weighted leaf absorptance, and photosynthetic performance

After 15 days of continuous irradiation under combined red and blue LED (LED_R+B_) or LD (LD_R+B_), leaf absorptance spectra (400–700 nm) were determined for all species (Fig. 5A). In tobacco, leaves grown under LD_R+B_ exhibited higher absorptance across most wavelengths compared with those grown under LED_R+B_, with particularly pronounced differences in the green and red regions. Consequently, spectrally weighted leaf absorptance (A_s_) was 13.2% higher under LD_R+B_ than under LED_R+B_ (Fig. 5B). In lettuce, higher absorptance under LD_R+B_ was mainly observed in the red region, resulting in a 9.5% increase in A_s_ compared with LED_R+B_. In Arabidopsis, LED_R+B_-grown leaves showed slightly higher absorptance in the green region, whereas LD_R+B_-grown leaves exhibited marginally higher absorptance in the red region. Nevertheless, A_s_ under LD_R+B_ was 2.5% greater than under LED_R+B_ (Fig. 5B). In both tobacco and lettuce, enhanced light absorptance under LD_R+B_ was accompanied by higher photosynthetic performance (Fig. 5C and D). The quantum efficiency of PSII [Y(II)] was significantly higher under LD_R+B_, increasing by 7.2% in tobacco and 6.9% in lettuce. Accordingly, electron transport rate (ETR) was also significantly higher under LD_R+B_, by 21.4% in tobacco and 17.1% in lettuce. In contrast, no significant differences in Y(II) or ETR were detected between light treatments in Arabidopsis.

**Figure 5.**
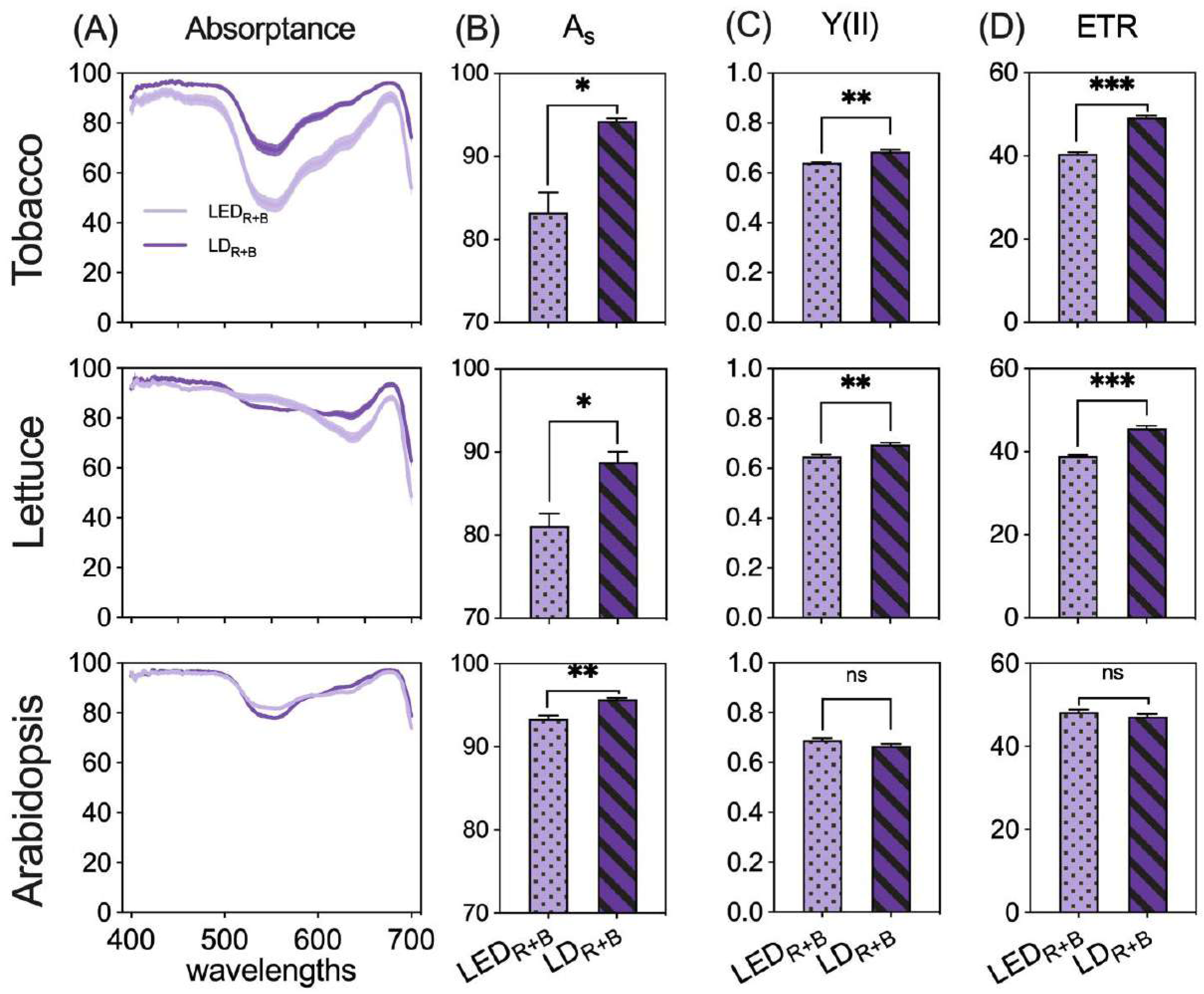
Leaf absorptance characteristics and Photosystem II performance of tobacco, lettuce, and Arabidopsis. (A) Leaf absorptance (%) spectra (400–700 nm). (B) Spectrally weighted leaf absorptance (A_s_; %) derived by integrating the leaf absorptance spectra (A) with the respective light source. (C) Effective quantum yield of PSII (Y(II)). (D) Electron transport rate (ETR; μmol m^-2^ s^-1^). In all panels, rows represent tobacco (top), lettuce (middle), and Arabidopsis (bottom). Data are presented as mean ± SE (n=4). Asterisks indicate significant differences between treatments ((* *P* < 0.05, ** *P* < 0.01, *** *P* < 0.001; *t* test). ns indicates no significant difference.

#### 3.2.2. Plant growth and morphology

Distinct differences in plant morphology were observed after 15 days of continuous combined red and blue irradiation (Fig. 6A). Across all species, plants grown under LD_R+B_ appeared visibly larger than those grown under LED_R+B_. Young tobacco leaves under LD_R+B_ displayed a greener appearance, whereas lettuce and Arabidopsis plants grown under LED_R+B_ exhibited more pronounced anthocyanin pigmentation. In addition, Arabidopsis under LED_R+B_ initiated bolting and flowering earlier than plants grown under LD_R+B_ (Fig. 6A). Quantitative analyses confirmed these observations (Fig. 6B). In tobacco, LD_R+B_ resulted in a 27.5% increase in shoot fresh weight and a 27.1% increase in total leaf area relative to LED_R+B_. These changes were accompanied by a significant increase in leaf angle (+10.8°) and a reduction in leaf mass per area (LMA). Lettuce showed similar trends, with LD_R+B_ increasing shoot fresh weight by 20.9% and total leaf area by 19.4%, along with a 16.6° increase in leaf angle and a 36.0% reduction in LMA. Arabidopsis exhibited the strongest response, with LD_R+B_ increasing shoot fresh weight and total leaf area by 84.9% and 84.5%, respectively. This was associated with a 20.0° increase in leaf angle and a 45.6% decrease in LMA. Shoot dry weight was comparable between light treatments, while the increases in shoot fresh weight under LD_R+B_ were accompanied by higher leaf water content (Supplementary Fig. 3).

**Figure 6.**
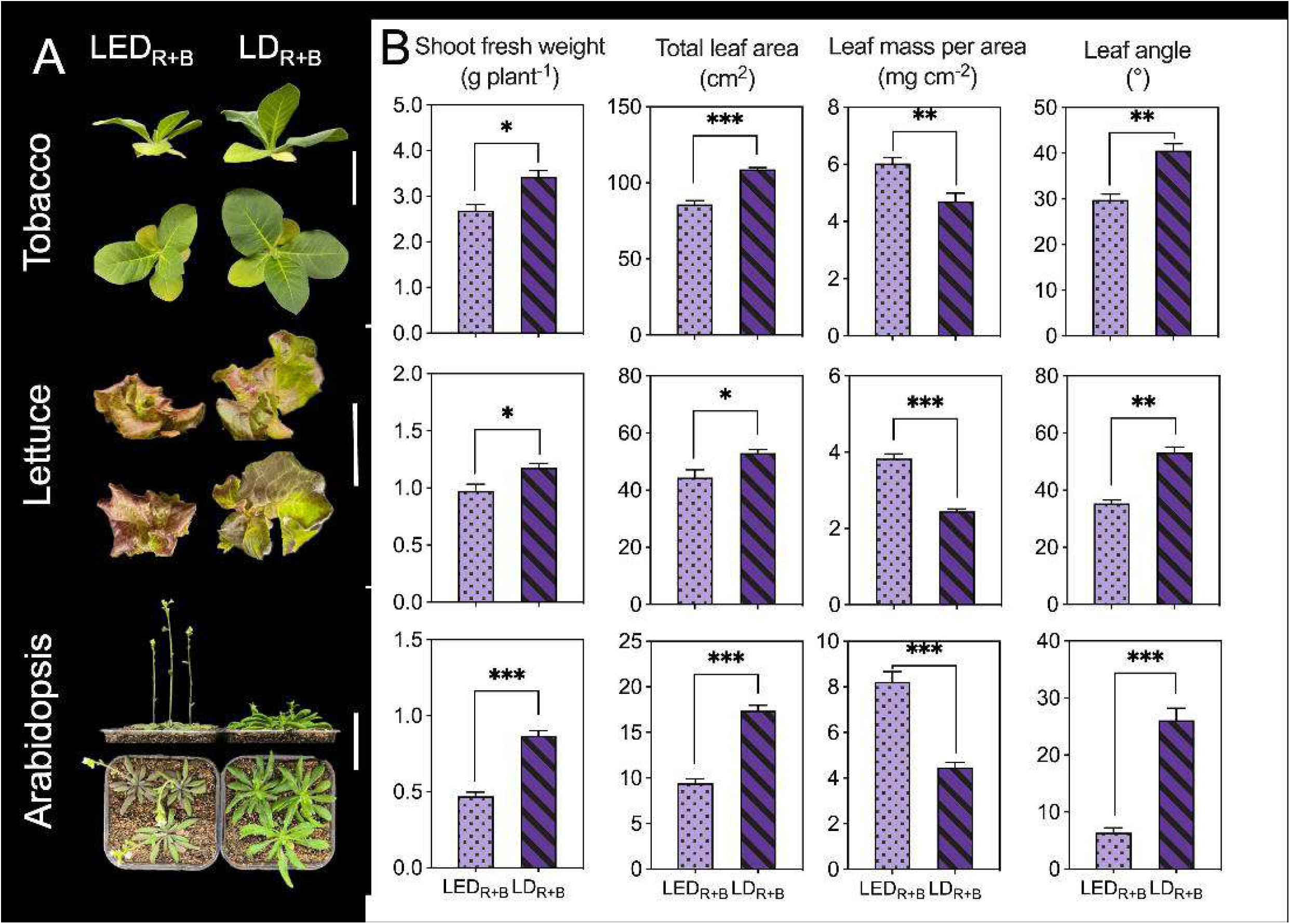
Growth and morphology of tobacco, lettuce, and Arabidopsis under continuous combined red and blue light. Plants were grown for 15 days under either a LED (LED_R+B_) or a LD (LD_R+B_) at of PPFD of 150 μmol·m□²·s□¹. (A) Representative images of plants (side and top views). The white scale bar in each picture represents 5 cm. (B) Shoot fresh weight, total leaf area, leaf mass per area, and leaf angle. Data are presented as the mean ± SE (n=4). Asterisks indicate significant differences between treatments (* *P* < 0.05, ** *P <* 0.01, *** *P* < 0.001; *t* test). ns indicates no significant difference.

#### 3.2.3 Pigment content

Representative leaf phenotypes of tobacco under LED_R+B_ and LD_R+B_ treatments are illustrated in Fig. 7A, revealing distinct differences in leaf size and coloration. Notably, the young fully expanded leaves of tobacco grown under LD_R+B_ were visibly larger and exhibited a deeper green color compared to those under LED_R+B_. These observations were supported by quantitative analysis of leaf pigments (Fig. 7B). Relative chlorophyll content (SPAD value) was significantly enhanced by the LD_R+B_ treatment across all three species. Specifically, compared to LED_R+B_, SPAD values under LD_R+B_ were 28.7%, 25.1%, and 17.5% higher in tobacco, lettuce, and Arabidopsis, respectively. In contrast, anthocyanin content (ACI value) followed an opposite trend (Fig. 7C). While anthocyanins were undetectable in tobacco under either light treatment, they were significantly lower in both lettuce and Arabidopsis under LD_R+B_, showing reductions of 53.7% and 59.5%, respectively, compared to LED_R+B_.

**Figure 7.**
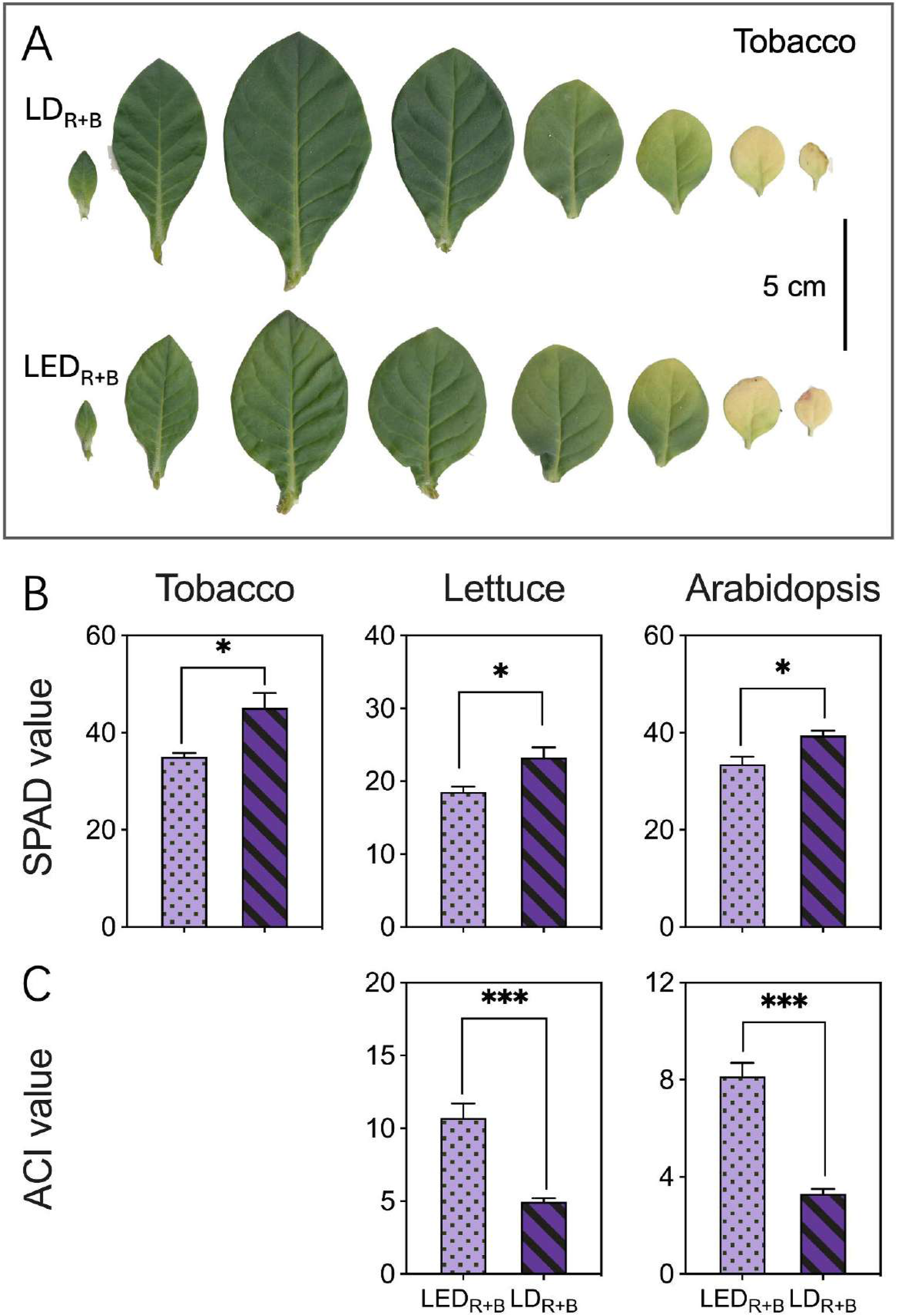
Leaf phenotype of tobacco and pigment content of tobacco, lettuce, and Arabidopsis under continuous combined red and blue light. Plants were grown under continuous light either from combined LEDR+B or LDR+B at a PPFD of 150 μmol·m□²·s□¹ for 15 days. (A) Leaf phenotypes of a representative tobacco plant from LED_R+B_ or LD_R+B_, illustrating differences in size and coloration between treatments. (B) Relative chlorophyll content (SPAD value) and (C) anthocyanin content (ACI value) for the three plant species. Data are presented as the mean ± SE (n=4). Asterisks indicate significant differences between treatments (* P < 0.05, *** P < 0.001; t test). N.D. indicates not detectable.

## 4. Discussion

Extensive research has used light-emitting diodes (LEDs) to optimize spectral recipes for indoor horticulture, largely focusing on peak wavelength composition to enhance productivity according to species and production objectives (Ohtake et al. 2018, 2021; Saengtharatip et al., 2021; Budavári et al., 2024). However, because LEDs emit relatively broad wavebands, the physiological consequences of spectral bandwidth itself, particularly when peak wavelength is identical, remain poorly understood. Laser diodes (LDs), which emit extremely narrow wavebands, provide an effective tool to address this gap. Building on our previous finding that monochromatic red LDs enhanced photosynthesis and growth relative to red LEDs (Li et al., 2025), the present study examined how spectral bandwidth influences plant responses under monochromatic blue and combined red and blue light. We show that even with identical peak wavelengths, differences in bandwidth substantially altered plant growth, canopy architecture, and senescence patterns. Under monochromatic blue light, blue LED promoted greater dry matter accumulation, whereas blue LD induced a more upright canopy and preferentially maintained chlorophyll content in lower leaves, indicating alleviated lower-canopy senescence (Figs. 3 and 4). Thus, bandwidth-dependent effects decoupled biomass accumulation from canopy maintenance within the same spectral color. When red and blue light were combined, LD lighting promoted steeper leaf angles, enhanced photosynthetic performance, and increased total leaf area, resulting in higher shoot fresh weight across species through coordinated canopy- and leaf-level responses (Figs. 5 and 6). Together, these results highlight spectral bandwidth as an important but underappreciated parameter in horticultural lighting control, capable of modulating plant architecture, senescence, and growth even when peak wavelength and light intensity are held constant.

### 4.1 Beyond peak wavelength: the critical role of blue light bandwidth in photosynthetic performance

Blue light is essential for photosynthesis, driving both energy capture and regulatory processes that enhance photosynthetic efficiency (Kochetova et al., 2022). However, its effectiveness varies considerably depending on spectral peak and bandwidth. Early studies by McCree (1972), using 25 nm spectral intervals, identified 450 nm as the most effective wavelength for CO□ assimilation across 22 growth chamber and 8 field plant species. Later, Inada (1976), with improved intervals (8.5–17 nm), shifted this peak to 435 nm across 33 species. In our study, LD 407 achieved significantly higher P_n_ than LD 450 despite both treatments having the same narrow bandwidth (Fig. 2A), suggesting that wavelength-specific effects can be obscured when spectral resolution is limited. Notably, despite sharing the same emission peak, LED 450 (44 nm bandwidth) outperformed LD 450 (7 nm bandwidth) (Fig. 2A), indicating that peak wavelength alone is insufficient to predict photosynthetic performance under blue light. A plausible explanation is that the broader spectrum of LED 450 supplied photons across multiple regions of high quantum yield, consistent with the absorption characteristics of photosynthetic pigments (Supplementary Fig. 1A). The absorption peak of chlorophyll a in the blue region is located at approximately 435 nm, while that of chlorophyll b is around 470 nm, both of which are covered by LED 450 but not by the narrow-band LD 450.

Despite differences in P_n_, stomatal conductance remained unchanged among treatments (Fig. 2B), implying that bandwidth-dependent variation in assimilation may be driven by non-stomatal processes. Accordingly, intrinsic water use efficiency mirrored the trend in P_n_ (Fig. 2C), supporting the idea that optimizing both peak wavelength and bandwidth can improve carbon gain per unit water loss in controlled environments (Lawson and Blatt, 2014; Katsuhama et al. 2025).

### 4.2 Blue LD displayed more vertical leaf angles, leading to enhanced light penetration and reduced the senescence of lower leaves

A plant factory with artificial lighting is a controlled production system designed to achieve high yields while maintaining product quality. However, high planting densities in these systems create light distribution challenges, particularly in lower canopy layers. The outer leaves beneath the dense canopy receive insufficient light, leading to accelerated senescence and requiring removal, which can result in yield losses of up to 10% (Zhang et al., 2015). Previous studies have proposed supplemental upward LED lighting to delay senescence in leafy vegetables and cut flowers (Zhang et al., 2015; Joshi et al., 2017; Saengtharatip et al., 2021; Yamori et al., 2021), but such approaches add infrastructural complexity.

Here, we demonstrate an alternative strategy that does not require modifying lighting geometry. Plants grown under LD_B_ (LD 450) exhibited more vertical leaf angles than those grown under LED_B_ (LED 450), improving light penetration into the lower canopy and delaying senescence, as indicated by higher chlorophyll status in lower leaves (Figs. 3 and 4). These results highlight a key practical distinction: under identical peak wavelength, broad-band blue LED enhanced biomass accumulation, whereas narrow-band blue LD promoted canopy architectures that better preserve lower leaves during dense cultivation.

The leaf angle divergence likely reflects differences in photoreceptor sensitivity to spectral distribution and spatial light patterns. The broad spectrum of LED_B_ overlaps more strongly with phototropin absorption peaks near 440 and 470 nm (Supplementary Fig. 1) and can promote leaf flattening under uniform blue light via phototropin signalling (Inoue et al., 2008). In contrast, the coherence of LD_B_ may generate micro-scale speckle and spatial heterogeneity (Pieczywek et al., 2024), which can be perceived through phototropin-linked directional growth pathways involving auxin transport, leading to gradual hyponasty (Legris et al., 2021). Although phytochrome absorbs in the blue region, calculated phytochrome photoequilibria were similar between treatments (Supplementary table 1) (Sager et al., 1998), supporting the view that the observed architectural differences primarily arise from blue-light–sensing pathways rather than phytochrome signaling.

### 4.3 Wavelength bandwidth of blue light influences photoprotection and secondary metabolite accumulation in plants

Blue photons not only drive photosynthesis but also regulate secondary metabolism. Monochromatic blue LED light has been shown to promote flavonoid synthesis, including anthocyanins (Landi et al., 2020; Bian et al., 2015). Consistent with this, lettuce plants exposed to both LED_B_ and LD_B_ accumulated anthocyanins (Supplementary Fig. 2A) (Anum et al., 2024a). Because anthocyanin accumulation serves as a photoprotective response by filtering excess light and/or scavenging reactive oxygen species (Landi et al., 2015), the stronger anthocyanin accumulation under LED_B_ suggests higher sustained light stress under broad-band blue illumination. This difference may be linked to canopy architecture. The more vertical leaf orientation under LD_B_ reduces light interception per unit leaf area and may distribute irradiance more evenly within the plant (Fig. 3), thereby alleviating overexcitation. In addition, LED_B_ rapidly induced higher nicotine accumulation in tobacco compared with LD_B_ (Supplementary Fig. 2B), indicating that bandwidth-dependent effects can also shape rapid metabolic adjustments (Steppuhn et al., 2004; Anum et al., 2024b). Together, these results suggest that the bandwidth of blue light influences not only carbon gain and architecture but also photoprotection and secondary metabolite responses.

### 4.4 LD_R+B_ light mitigates continuous light stress while enhancing light capture and whole-plant performance

Red and blue light are typically combined in horticultural lighting because they jointly regulate key processes including chlorophyll accumulation, stomatal opening, photosynthesis, photoprotective pigmentation, and quality-related secondary metabolism (Bantis et al., 2018). Building on our previous finding that monochromatic red LDs enhance photosynthesis and growth relative to red LEDs (Li et al., 2025), we tested whether combining red and blue in narrow-band form (LD_R+B_) improves whole-plant performance under a demanding continuous-light regime. Under 24-hour continuous illumination, plants grown under LD_R+B_ maintained a healthier physiological state than those grown under LED_R+B_, as indicated by higher chlorophyll status and lower anthocyanin accumulation (Fig. 7). Continuous light is known to induce chlorosis and necrosis (Velez-Ramirez et al., 2011), and the stronger photoprotective responses under LED_R+B_ are consistent with greater photo-oxidative pressure. Similar mitigation of stress responses by laser-based irradiation has been reported at the protein-expression level (Ooi et al., 2016). Structural acclimation under LED_R+B_ also aligned with stress exposure, including higher LMA, which has been associated with protection of the photosynthetic apparatus under excess light (Amiard et al., 2005; Xiao et al., 2016). Crucially, LD_R+B_ combined stress mitigation with traits that enhance whole-plant light acquisition. Leaves acclimated to LD_R+B_ exhibited higher spectrally weighted absorptance (A_s_) (Fig. 5B) and, in tobacco and lettuce, higher Y(II) and ETR (Fig. 5C and D), indicating improved photochemical performance when weighted by the growth spectra. At the canopy level, LD_R+B_ promoted more upright leaf orientation and larger total leaf area (Fig. 6), which together increase the effective interception of light at the plant level and reduce the fraction of senescing leaves within dense canopies. This coordinated suite of responses provides a mechanistic explanation for the consistent increase in shoot fresh weight observed under LD_R+B_ across species, integrating enhanced photosynthetic performance with improved whole-plant light interception and reduced senescence (Fig. 6).

An additional conceptual outcome is that LD lighting enables partial uncoupling of the classic “sun/shade leaf syndrome”. While sun leaves typically combine upright orientation with thicker leaves and high LMA, shade leaves are usually flatter with low LMA and larger area (Terashima and Hikosaka, 1995; Evans and Poorter, 2001; Kim et al., 2005; Zhen et al., 2022; Yang et al., 2023). In our study, LD_B_ and LD_R+B_ produced a distinctive phenotype that combines upright canopy structure with low LMA and expanded leaf area, demonstrating that spectral bandwidth can be used as a practical lever to more independently tune architecture, acclimation, and growth strategy for production goals.

## 5. Conclusion

In conclusion, this study demonstrates that spectral bandwidth, in addition to peak wavelength, is a critical parameter in horticultural lighting control. Using laser diodes as narrow-band light sources, we show that blue light with identical peak wavelength but different bandwidths induces distinct growth strategies: broad-band blue LEDs enhance dry matter accumulation, whereas narrow-band blue LDs promote a more upright canopy that enhances light penetration and suppresses the senescence of lower leaves. Thus, spectral bandwidth emerges as an effective lever for regulating canopy architecture and health in dense cultivation systems. Under combined red and blue light, narrow-band LD lighting consistently improved whole-plant performance. LD_R+B_ mitigated stress induced by 24-hour continuous illumination and promoted coordinated increases in photosynthetic performance, leaf expansion, and canopy structure, resulting in higher shoot fresh weight across species. Importantly, these gains were achieved through coordinated improvements in whole-plant light interception and reduced senescence, in addition to enhanced photosynthetic performance. Overall, precise control of spectral bandwidth enabled partial decoupling of traits traditionally associated with ‘sun’ and ‘shade’ leaf syndromes, allowing more flexible regulation of plant architecture and acclimation. These findings position laser diodes as a promising next-generation light source for precision horticulture, offering new opportunities to optimize canopy health, yield, and quality in indoor horticulture.

## Data availability statement

Supporting data can be requested by contacting the corresponding author.

## Figure captions

**Supplementary figure 1.**
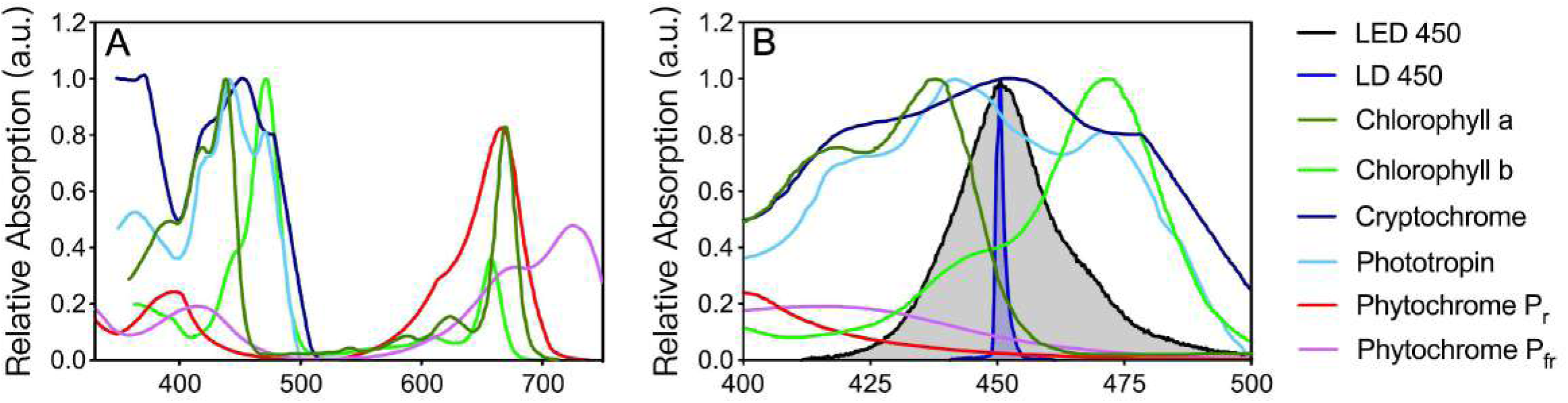
Blue light source spectra and the absorption spectra of key plant pigments and photoreceptors. (A) Normalized absorption spectra for major photosynthetic pigments (Chlorophyll a, Chlorophyll b) and key photoreceptors (Cryptochrome, Phototropin, Phytochrome P_r_, and Phytochrome P_fr_). (B) Spectral distributions of the LED 450 and LD 450 light sources, overlaid with the absorption spectra of pigments and photoreceptors active in the 400–500 nm range.

**Supplementary 2.**
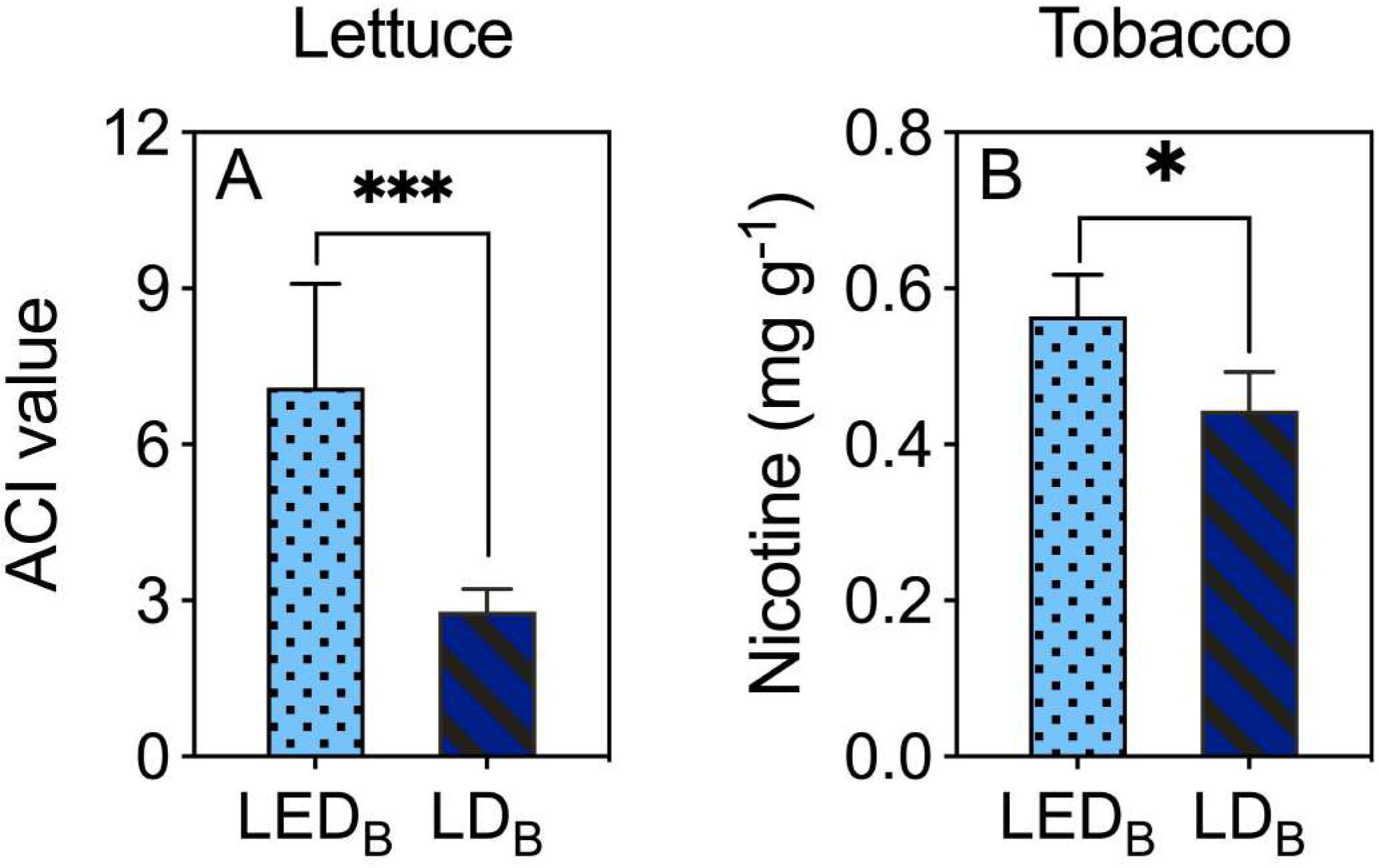
Secondary metabolite accumulation in tobacco and lettuce under monochromatic blue light. Treatments consisted of a monochromatic blue LED (LED_B_) or a blue LD (LD_B_), both with an identical peak wavelength at 450 nm, applied at a PPFD of 150 μmol·m□²·s□¹. (A) Anthocyanin accumulation (ACI value; leaf area based) in lettuce plants after 12 days of continuous irradiation. (B) Nicotine content of tobacco leaves after 8 hours of irradiation. Data are presented as the mean ± SE (n = 4). Asterisks indicate significant differences between treatments (* *P* < 0.05, *** *P* < 0.001; *t* test).

**Supplementary figure 3.**
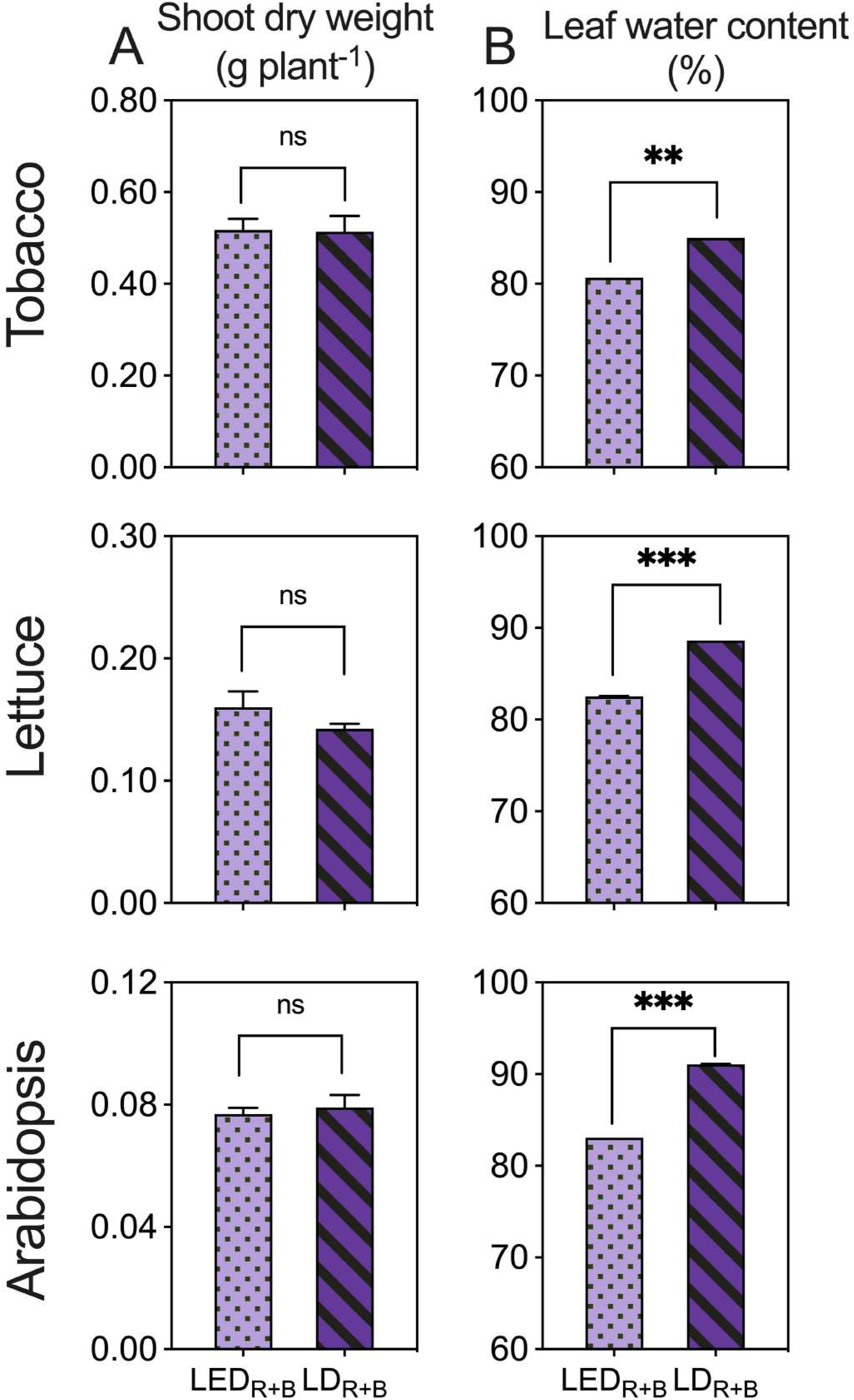
Shoot dry weight (A) and leaf water content (B) of tobacco, lettuce, and Arabidopsis under continuous combined red and blue light. Plants were grown for 15 days under either a LED (LED_R+B_) or a LD (LD_R+B_) at of PPFD of 150 μmol·m□²·s□¹. Data are presented as the mean ± SE (n=4). Asterisks indicate significant differences between treatments (** *P <* 0.01, *** *P* < 0.001; *t* test). ns indicates no significant difference.

**Supplementary Table 1.**
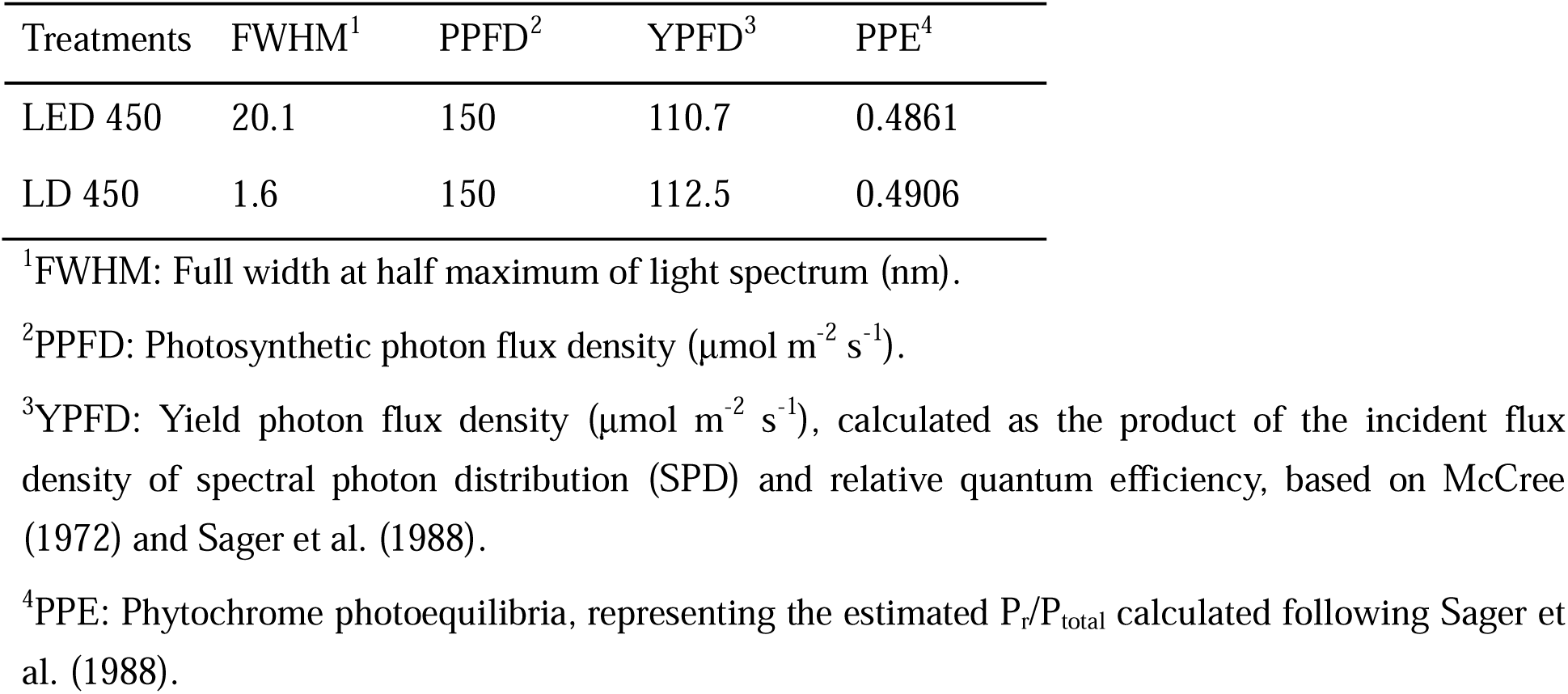
Spectral characteristics of LED 450 and LD 450.

